# Modular reconfiguration of an auditory-control brain network supports adaptive listening behavior

**DOI:** 10.1101/409797

**Authors:** Mohsen Alavash, Sarah Tune, Jonas Obleser

## Abstract

Speech comprehension in noisy, multi-talker situations poses a challenge. Human listeners differ substantially in the degree to which they adapt behaviorally and can listen successfully under such circumstances. How cortical networks embody this adaptation, particularly at the individual level, is currently unknown. We here explain this adaptation from reconfiguration of brain networks for a challenging listening task (i.e., a novel linguistic variant of the Posner paradigm with concurrent speech) in an age-varying sample of N = 49 healthy adults undergoing resting-state and task fMRI. We here provide evidence for the hypothesis that more successful listeners exhibit stronger task-specific reconfiguration, hence better adaptation, of brain networks. From rest to task, brain networks become reconfigured towards more localized cortical processing characterized by higher topological segregation. This reconfiguration is dominated by the functional division of an auditory and a cingulo-opercular module, and the emergence of a conjoined auditory and ventral attention module along bilateral middle and posterior temporal cortices. Supporting our hypothesis, the degree to which modularity of this fronto-temporal auditory-control network is increased relative to resting state predicts individuals’ listening success in states of divided and selective attention. Our findings elucidate how fine-tuned cortical communication dynamics shape selection and comprehension of speech. Our results highlight modularity of the auditory-control network as a key organizational principle in cortical implementation of auditory spatial attention in challenging listening situations.

**Significance Statement:** How do brain networks shape our listening behavior? We here develop and test the hypothesis that, during challenging listening situations, intrinsic brain networks are reconfigured to adapt to the listening demands, and thus to enable successful listening. We find that, relative to a task-free resting state, networks of the listening brain show higher segregation of temporal auditory, ventral attention, and frontal control regions known to be involved in speech processing, sound localization, and effortful listening. Importantly, the relative change in modularity of this auditory-control network predicts individuals’ listening success. Our findings shed light on how cortical communication dynamics tune selection and comprehension of speech in challenging listening situations, and suggest modularity as the network principle of auditory spatial attention.

## Introduction

Speech comprehension requires the fidelity of the auditory system, but it also hinges on higher-order cognitive abilities. In noisy, multi-talker listening situations speech comprehension turns into a challenging cognitive task for our brain. Under such circumstances, listening is often facilitated by a stronger engagement of auditory spatial attention (‘where’ to expect speech) and context-dependent semantic predictions (‘what’ to expect; Obleser et al., 2007; Davis et al., 2011; Wöstmann et al., 2016; Peelle, 2017; Dai et al., 2018). It has been long recognized that individuals differ substantially in utilizing these cognitive strategies in adaptation to challenging listening situations (e.g., Colflesh and Conway, 2007; Kidd et al., 2007; Tune et al., 2018). The cortical underpinning of this adaptation and the corresponding inter-individual variability are poorly understood, hindering future attempts to aid hearing or to rehabiliate problems in speech comprehension.

Functional neuroimaging has demonstrated that understanding degraded speech draws on cortical resources far beyond traditional perisylvian regions, involving cingulo-opercular, inferior frontal, and premotor cortices (Eckert et al., 2009; Obleser and Kotz, 2010; Adank, 2012; Wild et al., 2012; Erb et al., 2013; Vaden et al., 2013). While listening clearly marks a large-scale neural process shared across cortical nodes and networks (Price et al., 2005; Fedorenko and Thompson-Schill, 2014; Hagoort, 2014; Fuertinger et al., 2015; Chai et al., 2016) we do not know whether and how challenging speech comprehension relies on large-scale cortical networks (i.e., the functional connectome) and their adaptive reconfiguration.

The functional connectome is described by measuring correlated spontaneous brain responses using blood oxygen level dependent (BOLD) fMRI. When measured at rest, the result can be used to investigate whole-brain intrinsic networks (Raichle et al., 2000; Dosenbach et al., 2007; Power et al., 2011), which are relatively stable within individuals and reflect correlation patterns primarily determined by structural connectivity (Honey et al., 2007; Shen et al., 2015; Laumann et al., 2017). However, the make-up of resting state networks varies across individuals and undergoes subtle reconfigurations during the performance of specific tasks (Mueller et al., 2013; Cole et al., 2014; Finn et al., 2015; Geerligs et al., 2015; Gratton et al., 2016; Gordon et al., 2017a; Gordon et al., 2017b).

Accordingly, the variation from resting-state to task-specific brain networks has been proposed to predict individuals’ behavioral states or performance (Spadone et al., 2015; Cohen and D’Esposito, 2016; Gratton et al., 2016; Schultz and Cole, 2016; Bolt et al., 2017; Hearne et al., 2017). Interestingly, most of the variance in the configuration of brain networks arises from inter-individual variability rather than task contexts (Gratton et al., 2018). Here, we ask whether this variance contributes to the inter-individual variability in successful adaptation to a challenging listening situation.

Knowing that cortical systems supporting speech processing show correlated activity during rest (Tomasi and Volkow, 2012; Muller and Meyer, 2014; Tavor et al., 2016) we treat the whole-brain resting state network as the putative task network at its "idling" baseline (Figure 1A). As a listening challenge arises, different functional specializations for speech comprehension are conceivable across cortex depending on the task demands (Peelle, 2012). These functional specializations of speech comprehension can be studied at system-level within the framework of network segregation and integration using graph-theoretical connectomics (Sporns, 2013; Cohen and D’Esposito, 2016).

**Figure 1.**
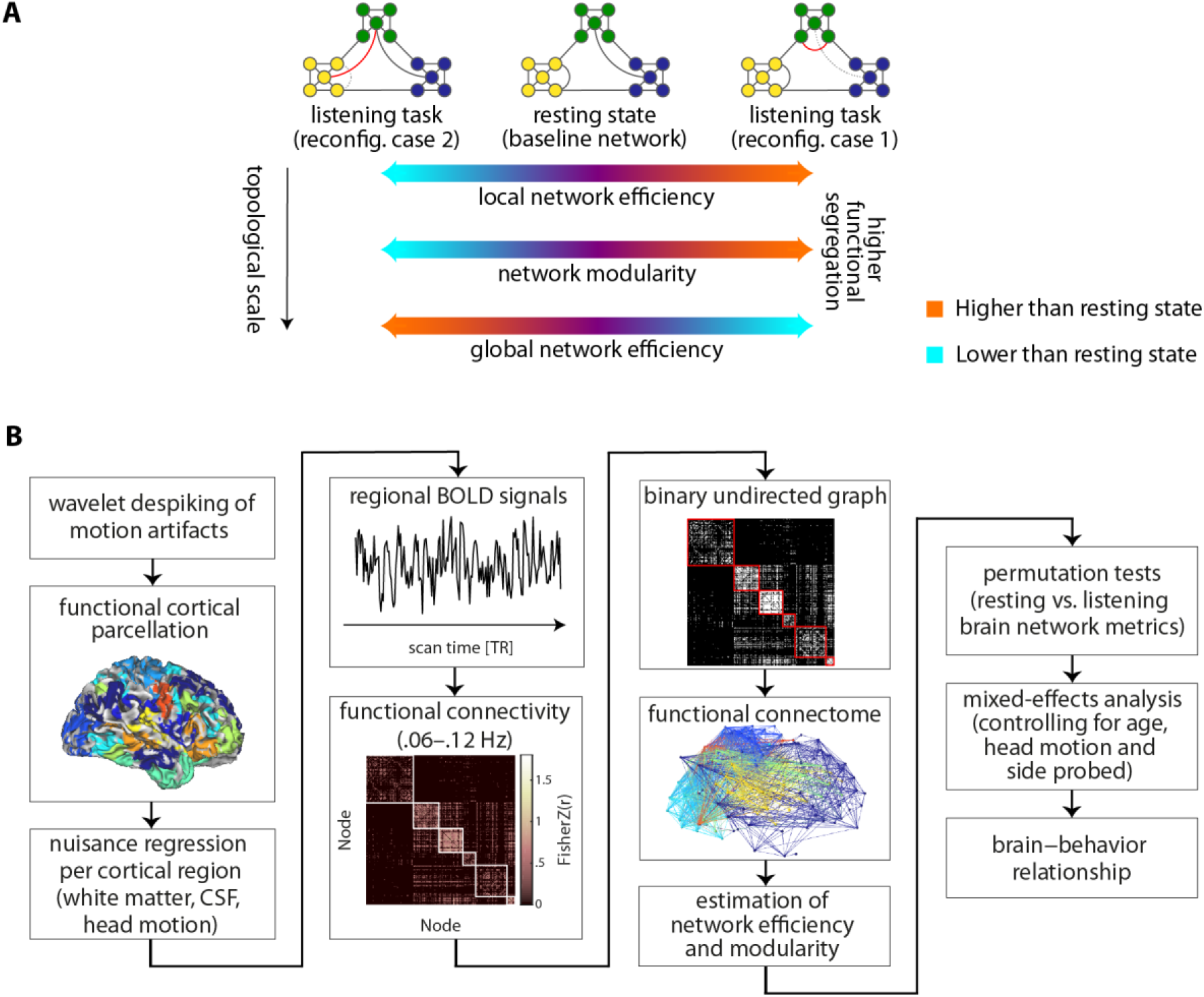
Methodological overview of the study. **(A)** Two possible network reconfigurations (case 1, case 2) characterized by a shift of the functional connectome towards either higher segregation (more localized cortical processing) or lower segregation (more distributed cortical processing} during a listening challenge. *Globalefficiency* is inversely related to the sum of shortest path lengths between every pair of nodes, indicating the capacity of a network for parallel processing. *Modularity* describes the segregation of nodes into relatively dense subsystems (here shown in distinct colors} which are sparsely interconnected. *Mean local efficiency* is equivalent to global efficiency computed on the direct neighbors of each node, which is then averaged over all nodes. Towards higher functional segregation, the hypothetical baseline network loses the shortcut between the blue and green module (dashed link). Instead, within the green module a new connection emerges (red link). Together, these form a segregated configuration tuned for a more localized cortical processing. **(B)** Analysis steps through which artifact-clean regional BOLD signals were averaged per cortical parcel. The results were used to construct graph-theoretical models of the functional connectome during resting state and the listening task, to ultimately investigate brain-behavior relationship using (generalized} linear mixed-effects analysis.

Accordingly, it is plausible to expect that attending to and processing local elements in the speech stream (DeWitt and Rauschecker, 2012; Vaden et al., 2013) would shift the topology of the functional connectome towards higher segregation (more localized cortical processing), while processing of larger units of speech (Lerner et al., 2011; de Heer et al., 2017) would lead to a reconfiguration towards lower segregation (more distributed cortical processing; Figure 1A).

As such, we here capitalize on the degree and direction of the functional connectome reconfiguration as a proxy for an individual’s successful adaptation, and thus speech comprehension success, in a challenging listening task (Figure 2A).

**Figure 2.**
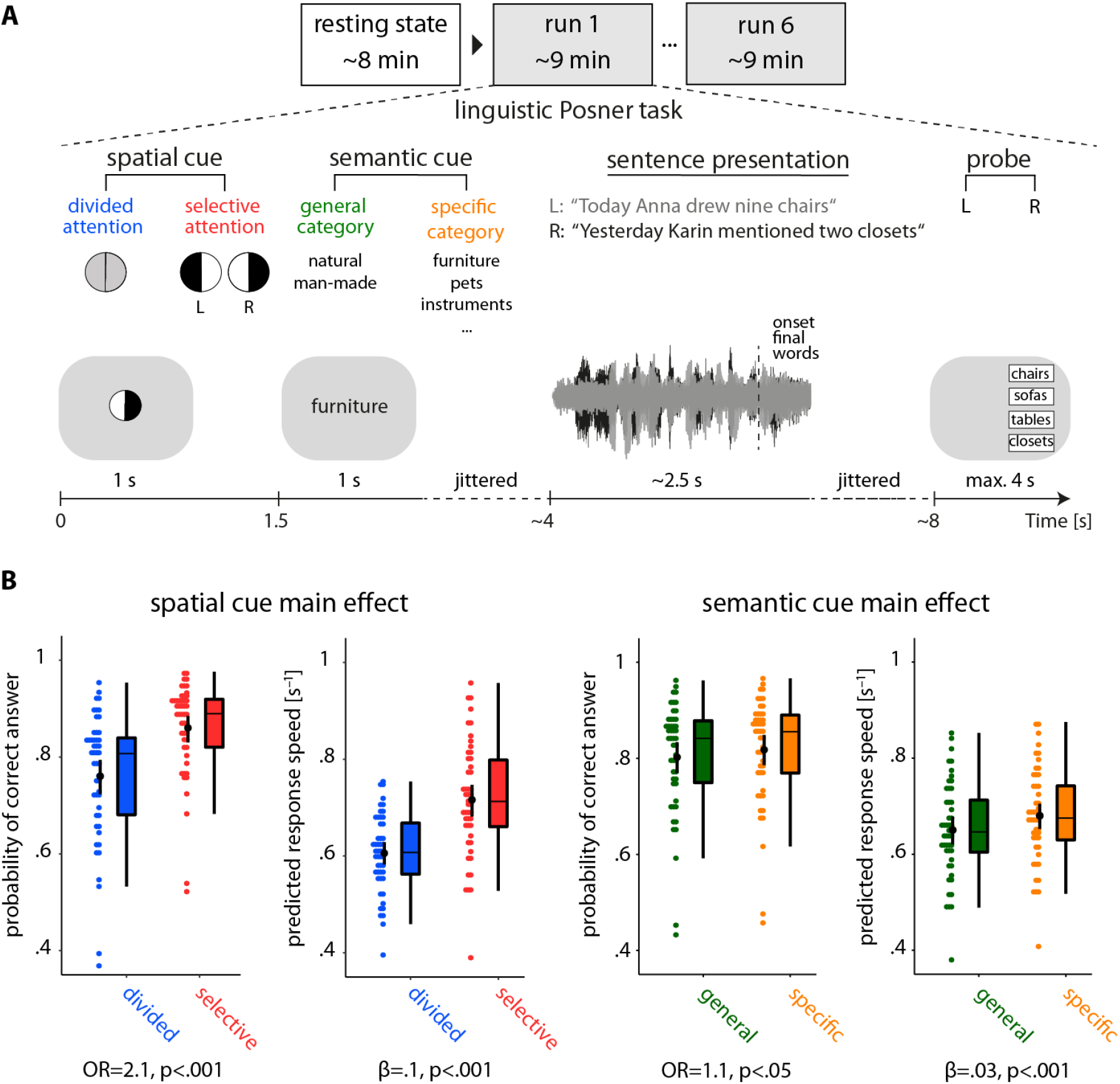
Listening task and individuals’ performance. **(A)** The linguistic Posner task with concurrent speech (i.e., cued speech comprehension). Participants listened to two competing, dichotically presented sentences. Each trial started with the visual presentation of a spatial cue. An informative cue provided information about the side (left ear vs. right ear) of to-be probed sentence-final word (invoking selective attention). An uninformative cue did not provide information about the side of to-be-probed sentence-final word (invoking divided attention). A semantic cue was visually presented indicating a general or a specific semantic category for both sentence-final words (allowing semantic predictions). The two sentences were presented dichotically along with a visual fixation cross. At the end of each trial a visual response array appeared on the left or right side of the screen with four word choices, asking the participant to identify the final word of the sentence presented to the left or right ear, depending on the side of the response array (L/R). **(B)** Predictions from linear mixed-effects models. Scattered data points (N=49) represent trial-averaged predictions derived from the model. Black points and vertical lines show mean± bootstrapped 95% Cl. OR: Odds ratio parameter estimate resulting from generalized linear mixed-effects models; *β*: slope parameter estimate resulting from general linear mixed-effects models.

## Results

We measured brain activity using fMRI in an age-varying sample of healthy adults (*N=*49; 19-69 yrs) and constructed large-scale cortical network models (Figure 1B) during rest and while individuals performed a novel linguistic Posner task with concurrent speech. The task presented participants with two competing, dichotic sentences uttered by the same female speaker (Figure 2A). Participants were probed on the last word (i.e., target) of one of these two sentences. Crucially, two visual cues were presented up-front: First, a spatial-attention cue either indicated the to-be-probed side, thus invoking selective attention, or it was uninformative, thus invoking divided attention. The second cue specified the semantic category of the sentence-final words either very generally or more specifically, which allowed for more or less precise semantic prediction of the upcoming target word. Using (generalized) linear mixed-effects models, we examined the influence of the listening cues and brain network reconfiguration on listening success, controlling for individuals’ age, head motion, and the probed side (Figure 1B).

### Informative spatial and semantic cues facilitate speech comprehension

The analysis of listening performance using linear mixed-effects models revealed an overall behavioral benefit from more informative cues. As expected, participants performed more accurately and faster under selective attention as compared to divided attention (accuracy: odds ratios (OR)= 2.1, p < 0.001; response speed: β= 0.1, p < 0.001; Figure 2B, first two panels). Moreover, participants performed more accurately and faster when they were cued to the specific semantic category of the target word as compared to a general category (accuracy: OR= 0.1, p = 0.02; response speed: β= 0.03, p < 0.01; Figu re 2B, last two panels). Participants benefitted notably more from the informative spatial cue than from the informative (i.e., specific) semantic cue. We did not find any evidence of an interactive effect of the two cues on behavior (likelihood ratio tests, accuracy: *χ^2^*=0.005, p = 0.9; response speed: *χ^2^* = 2.5, p = 0.1).

Lending validity to our results, younger participants performed more accurately and faster than older participants, as expected (main effect of age; accuracy: OR= 0.63, p < 0.0001; response speed: β= −0.03, p<0.001). Furthermore, as to be expected from the well-established right-ear advantage for linguistic materials (Hugdahl, 2003; Passow et al., 2013), participants were faster when probed on the right compared to the left ear (main effect of probe: β= 0.018, p < 0.01; see Table S1 for all model terms and estimates).

### Higher segregation of the whole-brain network during cued speech comprehension

A main question in the present study was whether and how resting state brain networks are reconfigured when listening challenges arise. We compared functional connectivity and three key topological network metrics between resting state and the listening task (see Supplemental Information for the definitions of brain metrics). We expected that resting state brain networks would functionally reshape and shift either towards higher segregation (more localized cortical processing) or lower segregation (more distributed cortical processing) during the listening task (Figure 1A). Our network comparison revealed how the whole-brain network reconfigured in adaptation to cued speech comprehension (Figure 3).

**Figure 3.**
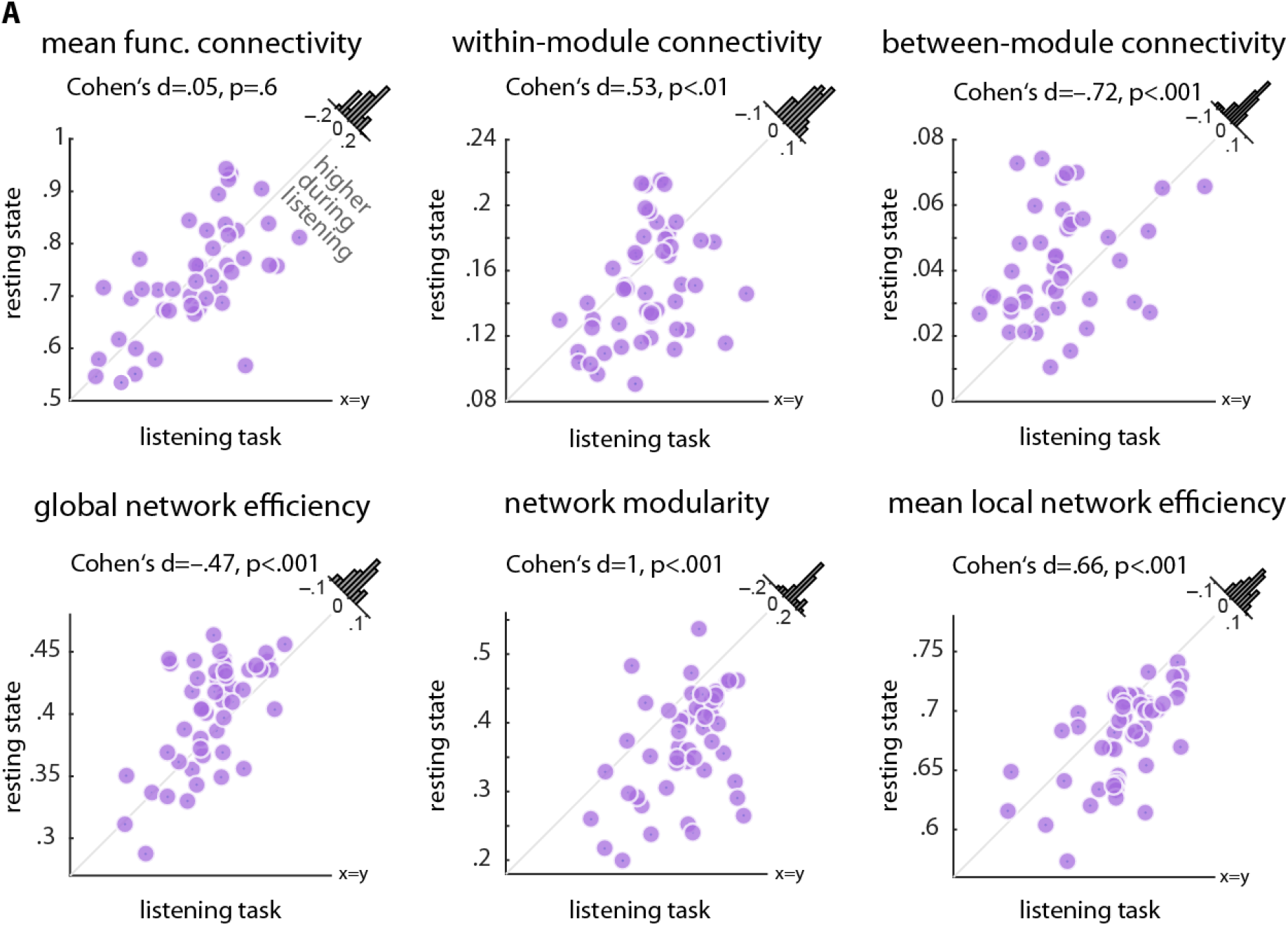
Alterations in whole-brain network metrics during the listening task relative to resting state. Functional segregation was significantly increased from rest to task. This was manifested in higher network modularity, within-module connectivity, and local network efficiency, but lower between-module connectivity and global network efficiency. Histograms show the distribution of the change (task minus rest) of the network metrics across all 49 participants.

First, while mean within-module functional connectivity of the whole-brain network showed a significant increase from rest to task (Cohen’s d = 0.53, p < 0.01; Figure 3, first row, middle panel), between-module connectivity showed the opposite effect (Cohen’s d = −0.72, p < 0.001; Figure 3, first row, last panel). Importantly, resting state and the listening-task networks did not differ in their overall mean functional connectivity (p = 0.6; Figure 3, first row, first panel), which emphasizes the importance of modular reconfiguration (i.e., change in network topology) in brain network adaptation in contrast to a mere alteration in overall functional connectivity (change in correlation strengths).

Second, during the listening task, functional segregation of the whole-brain network increased relative to resting state. This effect was consistently observed on the local, intermediate and global scales of network topology, a reconfiguration regime in the direction of the hypothetical case 1 depicted in Figure 1A.

More specifically, global network efficiency––a metric which has been proposed to measure the capacity of brain networks for parallel processing (Bullmore and Sporns, 2009)––decreased from rest to task (Cohen’s d = −0.47, p < 0.001; Figure 3, second row, first panel). In addition, modularity of brain networks––a metric related to the decomposability of a network into subsystems (Rubinov and Sporns, 2010)––increased under the listening task in contrast to resting state (Cohen’s d = 1, p < 0.001; Figure 3, second row, middle panel). Across participants, network modularity was higher in 43 out of 49 individuals, with a considerable degree of inter-individual variability in its magnitude. This result complements the significant changes in within- and between-module connectivity described earlier.

Moreover, the whole-brain network during the task was characterized by an increase in its local efficiency relative to resting state (Cohen’s d = 0.66, p < 0.001; Figure 3, second row, last panel). Knowing that local efficiency is inversely related to the topological distance between nodes within local cliques or clusters (Rubinov and Sporns, 2010), this effect is another indication of a reconfiguration towards higher functional segregation, reflecting more localized cortical processing. The above results were robust with respect to the choice of network density (Figure S1). Similar results were found when brain graphs were constructed (and compared to rest) separately per task block (Figure S2), which rules out any systematic difference between resting state and task due to the difference in scan duration (i.e., duration of each block∼ duration of resting state).

In an additional analysis, we investigated the correlation of brain network measures between resting state and listening-task state. Reassuringly, the brain network measures were all significantly positively correlated across participants (Figure S3), supporting the idea that state- and trait-like determinants both contribute to individuals’ brain network configurations (Geerligs et al., 2015).

We next investigated whether reconfiguration of the whole-brain networks from rest to task can account for the inter-individual variability in listening success. To this end, using the same linear mixed-effects model that revealed the beneficial effects of the listening cues on behavior, we tested whether the change in each whole-brain network metric (i.e., task minus rest) as well as its modulation by the different cue conditions could predict a listener’s speech comprehension success.

We found a significant interaction between the spatial cue and the change in individuals’ whole-brain network modularity in predicting accuracy (OR= 1.13, p < 0.01). This interaction was driven by a weak positive relationship of modularity and accuracy during selective attention trials (OR= 1.17, p = 0.13) and the absence of such a relationship during divided attention trials (OR = 1.02, p = 0.8; see Table S1 and S2 for all model terms and estimates). This result provides initial evidence that, at the whole-brain level, higher network modularity relative to resting state coincides with successful implementation of selective attention to speech. Notably, when changes in other network measures were included in the model (i.e., change in local or global network efficiency instead of change in network modularity), no significant relationships were found. Besides, the effect of the semantic cue on behavior was not significantly modulated by change in whole-brain network topology. More precisely, adding an interaction term between the semantic cue and change in brain network modularity did not significantly improve the model fit (likelihood ration tests; accuracy: *χ* ^2^ = 2.6, p = 0.1; response speed: *χ* ^2^ = 3.8, p = 0.06). There was also no significant interaction between age and change in network modularity in predicting behavior (likelihood ratio tests; accuracy: *χ* ^2^ = 1, p = 0.3; response speed: *χ* ^2^ = 3.1, p = 0.08).

In the following sections, we will explore the cortical implementation of the topological reconfiguration outlined above, first at the whole-brain level and then on a sub-network level, in more detail. In the final section of the results, we will illustrate the behavioral relevance of these reconfigurations at individual level.

### Reconfiguration of the whole-brain network in adaptation to cued speech comprehension

Figure 4 provides a comprehensive overview of group-level brain networks, functional connectivity maps and the corresponding connectograms (circular diagrams) during resting state (Figure 4A) and during the listening task (Figure 4C). In Figure 4, cortical regions defined based on Gordon et al. (2016) parcellation are grouped and color-coded according to their module membership determined using the graph-theoretical consensus community detection algorithm (Blondel et al., 2008; Rubinov and Sporns, 2010; see Supplemental Information for details).

**Figure 4.**
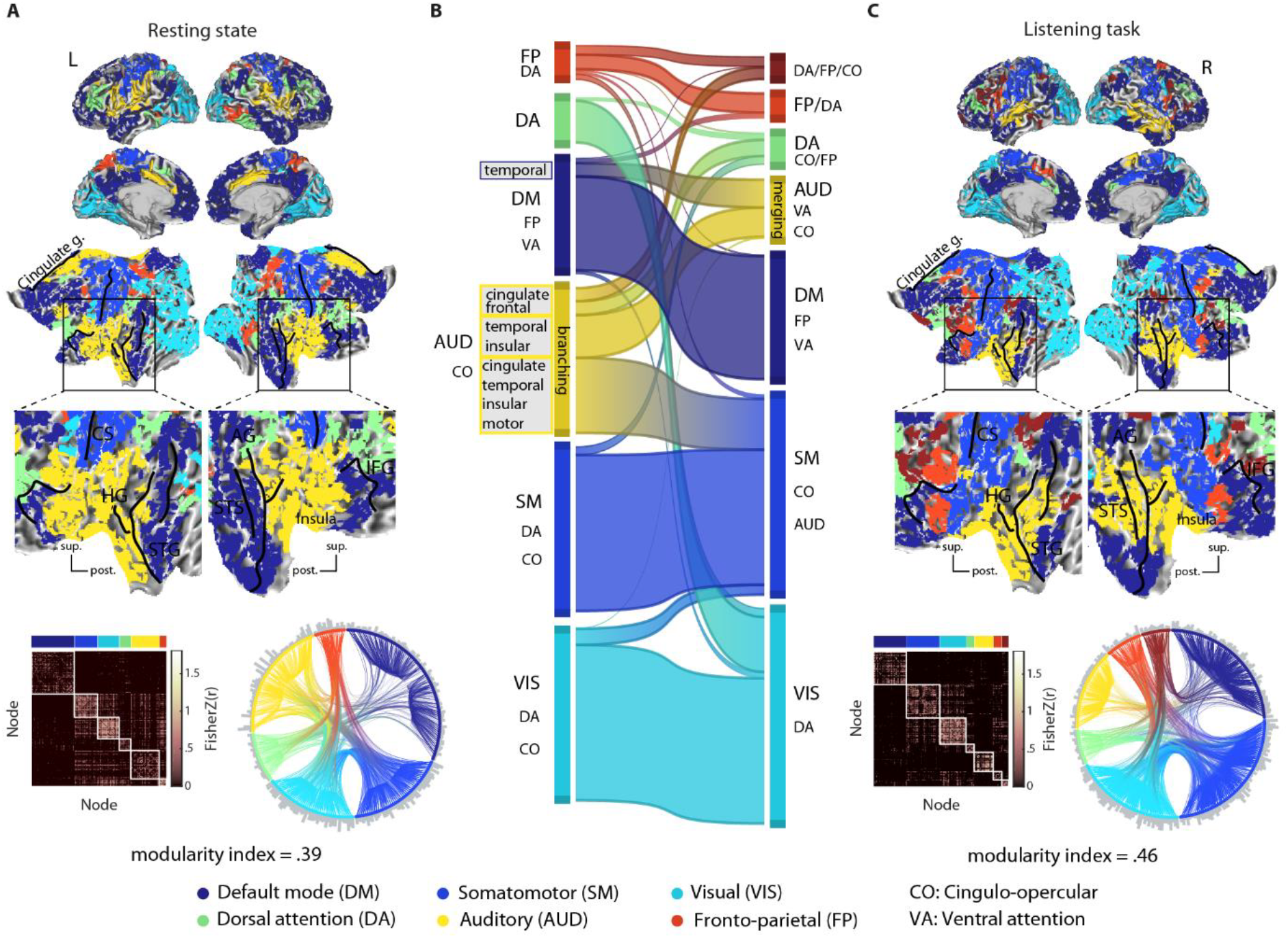
Modular reconfiguration of the whole-brain network in adaptation to the listening task. **(A)** The whole-brain resting state network decomposed into six distinct modules shown in different colors. The network modules are visualized on the cortical surface, within the functional connectivity matrix, and the connectogram using a consistent color scheme. Modules are functionally identified according to their node labels as in Gordon et al. (2016). Group-level modularity partition and the corresponding modularity index were obtained using graph-theoretical consensus community detection. Grey peripheral bars around the connectograms indicate the number of connections per node. **(B)** Flow diagram illustrating the reconfiguration of brain network modules from resting state (left) in adaptation to the listening task (right). Modules shown in separate vertical boxes at the left and right side are sorted from bottom to top according to the total PageRank of the nodes they contain, and their heights correspond to their connection densities. The streamlines illustrate how nodes belonging to a given module during resting state change their module membership during the listening task. **(C)** Whole-brain network modules during the listening task. The network construction and visualization scheme is identical to (A). *CS:* central sulcus; *HG:* Heschl’s gyrus; *IFG:* inferior frontal gyrus; *STG/S:* superior temporal gyrus/sulcus; *sup.:* superior; *post.:* posterior.

When applied to the resting state data in our sample, the algorithm decomposed the group-averaged whole-brain network into six modules (Figure 4A). The network module with the highest number of nodes largely overlapped with the known default mode network (Figure 4A, dark blue). At the center of our investigation lied the second largest module, comprising mostly auditory nodes and a number of cingulo-opercular nodes (Figure 4A, yellow). For simplicity, we will refer to this module as the auditory (AUD) module. Further network modules included (in the order of size) a module made up of visual nodes (Figure 4A, cyan), one encompassing mostly somatomotor nodes (Figure 4A, light blue), as well as a dorsal attention module (Figure 4A, green) and a fronto-parietal module (Figure 4A, red).

Figure 4B shows how the configuration of whole-brain network changed from rest to task. This network flow diagram illustrates whether and how cortical nodes belonging to a given network module during resting state (Figure 4B, left) changed their module membership during the listening task (Figure 4B, right). The streamlines tell us how the nodal composition of network modules changed, indicating a functional reconfiguration. According to the streamlines, the auditory module showed the most prominent reconfiguration (yellow module; AUD), while the other modules underwent only moderate reconfiguration.

This reconfiguration of the auditory module can be summarized by two dominant nodal alterations. On the one hand, a nodal "branching" arises: A number of cingulate, frontal, temporal and insular nodes change their module membership in adaptation to the listening task. On the other hand, a group of temporal and insular nodes merge with other temporal nodes from the default mode network module (Figure 4B, left side, dark blue) and form a new common module during the listening task (Figure 4B, right side, yellow). Note that nodal merging should not be mistaken with functional integration. In a graph-theoretical sense, higher functional integration would be reflected by stronger between-module connectivity which is opposite to what is found in the present study (Figure 3).

Interestingly, this latter group of temporal nodes includes posterior temporal nodes that are functionally associated with the ventral attention network (Corbetta et al., 2008; Power et al., 2011; Gordon et al., 2016). Considering the cortical surface maps, these alterations occur at the vicinity of the middle and posterior portion of the superior temporal lobes (note dark blue regions in the right superior temporal sulcus/gyrus in resting-state maps which turn yellow in the listening-state maps).

In line with the changes described above, the reconfiguration of the auditory module is also observed as a change in the connection patterns shown by the connectograms (Figure 4A and C, circular diagrams). Relative to rest, tuning in to the listening task condenses the auditory module, and its nodes have fewer connections to other modules. In addition, a group of cingulate and frontal nodes from the auditory module at rest merge with several nodes from the fronto-parietal/dorsal attention module, and form a new module during the listening task (Figure 4B, right side, dark brown; see also Figure 4C).

Taken together, we have observed a reconfiguration of the whole-brain network that is dominated by modular branching and merging across auditory, cingulo-opercular and ventral attention nodes (Figure 4B).

### Reconfiguration of a fronto-temporal auditory-control network in adaptation to cued speech comprehension

As outlined above, we had observed two prominent reconfigurations described in Figure 4B, namely nodal branching and nodal merging. Accordingly, we identified the cortical regions involved in these nodal alterations based on the underlying functional parcellation (Gordon et al., 2016). The identified nodes include auditory (AUD), ventral attention (VA), and cingulo-opercular (CO) regions. We will refer to these fronto-temporal cortical regions collectively as the auditory-control network. This network encompasses 86 nodes (out of 333 whole-brain nodes). According to Figure 4B, this network is in fact a conjunction across all auditory, ventral attention and cingulo-opercular nodes, irrespective of their module membership. For the purpose of a more transparent illustration, the cortical map of the auditory-control network is visualized in Figure 5A.

**Figure 5.**
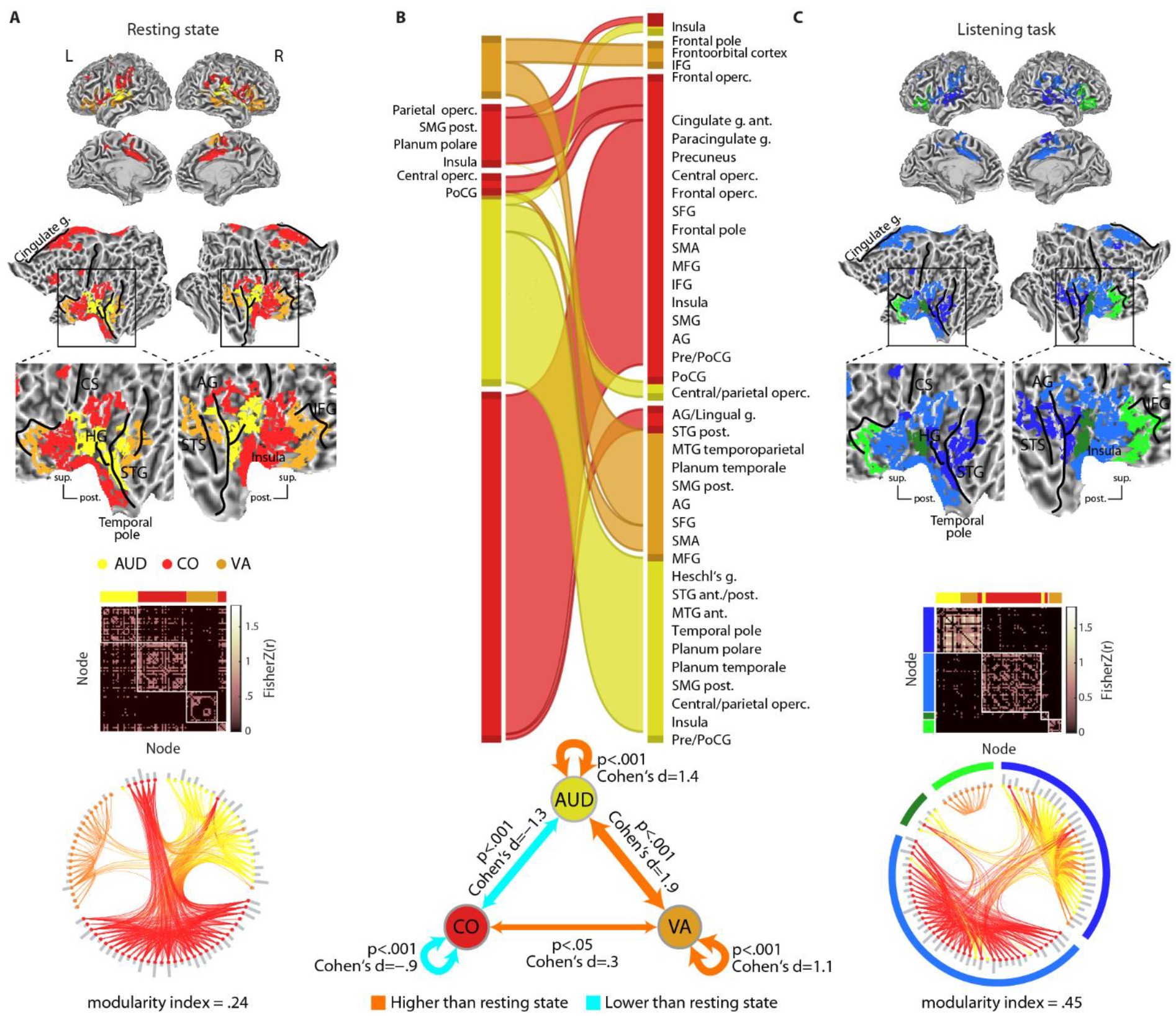
Modular reconfiguration of the fronto-temporal auditory-control network in adaptation to the listening task. **(A)** The auditory-control network. Cortical regions across the resting state fronto-temporal map are functionally identified and color-coded according to their node labels as in Gordon et al. (2016). This network is decomposed into four distinct modules shown within the group-level functional connectivity matrix and the connectogram (circular diagram). Group-level modularity partition and the corresponding modularity index were obtained using graph-theoretical consensus community detection. Grey peripheral bars around the connectograms indicate the number of connections per node. **(B)** *Top* Flow diagram illustrating the reconfiguration of the auditory-control network from resting state (left) to the listening task (right). Modules shown in separate vertical boxes at the left and right side are sorted from bottom to top according to the total PageRank of the nodes they contain, and their heights correspond to their connection densities. The streamlines illustrate how nodes belonging to a given module during resting state change their module membership during the listening task. *Bottom* Alteration in functional connectivity within the auditory-control network complements the topological reconfiguration illustrated by the flow diagram **(C)** Modules of the auditory-control network during the listening task. The network construction and visualization scheme is identical to (A). Since auditory and ventral attention nodes are merged (yellow and orange nodes), an additional green-blue color-coding is used for a clearer illustration of modules. *AG:* angular gyrus; *CS:* central sulcus; *HG:* Heschl’s gyrus; *IFG:* inferior frontal gyrus; *MFG:* middle frontal gyrus; *Operc.:* operculum; *Pre/PostCG:* Pre/post central gyrus; *SFG:* superior frontal gyrus; *SMA:* supplementary motor area; *SMG:* suparmarginal gyrus; *STG/5:* superior temporal gyrus/sulcus; *g.:* gyrus; *ant.:* anterior; *sup.:* superior; *post.:* posterior.

Similar to the whole-brain analysis (Figure 4), the auditory-control network can be decomposed into modules by applying the consensus community detection algorithm again, but this time to the graph encompassing the 86 nodes. The result is visualized in Figure SA by grouping the nodes within the group-level functional connectivity matrix and the corresponding connectogram. During resting state, the consensus community detection algorithm had decomposed the auditory-control network into four distinct modules that correspond well with their functional differentiation as in Gordon et al. (2016). That is, auditory, ventral attention, and cingulo-opercular nodes form their own distinct modules with almost no overlap.

During the listening task, however, the configuration of the auditory-control network changed (Figure 5B, top). According to the streamlines, the majority of auditory nodes (yellow) and posterior ventral attention nodes (orange) merge to form a common network module during the task (Figure 5B, right, first module from bottom). This reconfiguration co-occurred with an increase in functional connectivity within the auditory nodes, ventral attention nodes, as well as between auditory and ventral attention nodes (Figure 5B, bottom; see also Figure S4). The cortical and connectivity maps of the AUD-VA module are shown in Figure 5C (dark blue module).

This network module emerging during the listening task involves cortical regions mostly in the vicinity of the bilateral superior temporal sulcus (STS), Heschl’s gyrus (HG) as well as the right angular gyrus (AG). Note also the increase in the functional connectivity across these nodes (Figure 5C, connectivity matrix, dark blue module) and the increase in the number of connections within this module (Figure 5C, connectogram, grey bars inside dark blue module; all relative to resting state). Collectively, this reconfiguration indicates a stronger functional cross-talk between auditory and posterior ventral attention regions during the listening task as compared to resting state.

In contrast, frontal ventral attention nodes do not merge with other nodes, and remain connected within a separate module both during resting state and task (Figure 5B, right, second module from top). These nodes overlap with cortical regions at the vicinity of inferior frontal gyrus (IFG; Figure 5C, light green module). Moreover, the majority of cingulo-opercular nodes (red) are grouped together during the listening task (Figure 5C, light blue module). This co-occurs with a decrease in functional connectivity within the cingulo-opercular nodes, as well as between auditory and cingulo-opercular nodes (Figure 5B, bottom; see also Figure S4).This indicates higher functional segregation within the cingulo-opercular regions during the listening challenge relative to resting state.

### Higher modularity of the auditory-control network predicts speech comprehension success

To quantitatively assess the reconfiguration of the auditory-control network described above, we compared functional connectivity and topological metrics derived from this network between resting state and the listening task (Figure 6A).

**Figure 6.**
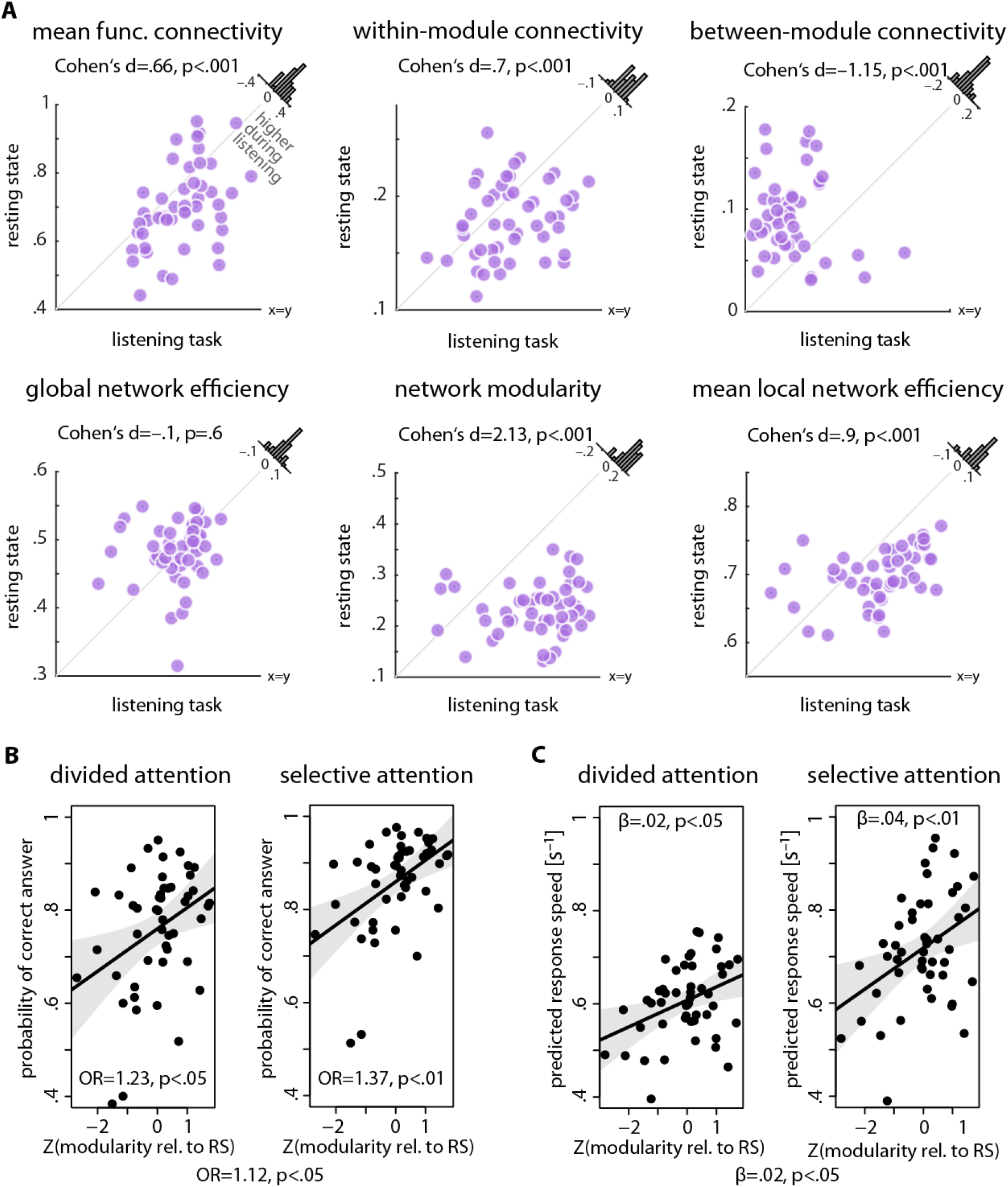
Brain network metrics derived from the fronto-temporal auditory-control network and prediction of listening success from modularity of this network. **(A)** Functional segregation within the auditoy-control network was significantly increased during the listening task relative to resting state. This was manifested in higher network modularity, within-module connectivity, and local network efficiency, but lower between-module connectivity. Histograms show the distribution of the change (task minus rest) of the network metrics across all 49 participants. **(B/C)** Interaction of spatial cue and change in modularity of the auditory-control network. The data points represent trial-averaged predictions derived from the (generalized) linear mixed-effects model. Solid lines indicate linear regression fit to the trial-averaged data. Shaded area shows two-sided parametric 95% Cl. *RS:* resting state; *OR:* Odds ratio parameter estimate resulting from generalized linear mixed-effects models; /3: slope parameter estimate resulting from general linear mixed-effects models.

First, while mean overall connectivity and within-module connectivity of this network significantly increased from rest to task (mean connectivity: Cohen’s d = 0.66, p < 0.001; within-module connectivity: Cohen’s d = 0.7, p < 0.001), between-module connectivity showed the opposite effect (Cohen’s d = −1.15, p < 0.001; Figure 6A, first row).

Second, functional segregation of the auditory-control network was significantly increased from rest to task, as manifested by higher degree of modularity and mean local efficiency (modularity: Cohen’s d = 2.13, p < 0.001; local efficiency: Cohen’s d = 0.9, p<0.001; Figure 6A, second row).

Critically, (generalized) linear mixed-effects analyses revealed significant interactions of the spatial cue and change in modularity of the auditory-control network for both behavioral dependent measures. More precisely, the change in modularity of this network showed positive correlations with accuracy (Figure 6B) and response speed (Figure 6C) under both divided and selective attention trials. However, these brain-behavior correlations were stronger under selective (accuracy: OR= 1.37, p < 0.01; response speed:β = 0.04, p < 0.01) than under divided attention trials (accuracy: OR= 1.23, p < 0.05; response speed:β = 0.02, p < 0.05; See Table S3 for all model terms and estimates). We did not find any evidence for an interaction between the semantic cue and change in modularity of the auditory-control network in predicting behavior (likelihood ratio tests; accuracy: *χ^2^=* 0.3, p = 0.6; response speed: *χ^2^* = 0.4, p = 0.5). There was also no significant interaction between age and change in modularity of the auditory-control network in predicting behavior (accuracy: *χ ^2^ =* 0.15, p = 0.7; response speed:*χ* ^2^ = 0.4, p = 0.5).

Moreover, an additional linear mixed-effects model with resting state modularity and task modularity as two separate regressors revealed a significant main effect of the auditory-control network modularity during the task (but not rest) in predicting accuracy (OR = 1.35, p < 0.001). Additionally, there was a significant interaction between modularity of the auditory-control network during the task with the spatial cue in predicting response speed (β = 0.01, p < 0.05; see Table S4 for all model terms and estimates).

Apart from the nodal branching and merging in the auditory module, other reconfigurations, although less profound, can also be identified (see Figure 4B). These include modular reconfiguration of sub-networks encompassing (1) visual, dorsal attention and cingulo-opercular nodes; (2) somato-motor, cingulo-opercular and auditory nodes; (3) default-mode, fronto-parietal and ventral attention nodes; (4) dorsal-attention, fronto-parietal and cingulo-opercular. To investigate whether these reconfigurations predict listening behavior, we performed analyses analogous to the one undertaken for the auditory-control network. More precisely, using four separate linear mixed-effects models, we included the change (i.e., task minus rest) in network modularity of each of the four aforementioned sub-networks. None of these models were able to predict individual listening behavior (accuracy or response speed). Thus, the prediction of speech comprehension success from modular reconfiguration of brain sub-networks was specific to the fronto-temporal auditory-control network (Figure 6B).

## Discussion

We here have first shown how resting state brain networks reshape in adaptation to a challenging listening task. This reconfiguration predominantly emerges from a fronto-temporal auditory-control network which shows higher modularity and local efficiency during the listening task, shifting the whole-brain connectome towards higher segregation (more localized cortical processing). Second, the degree to which modularity of this auditory-control network increases relative to its resting state baseline predicts an individual’s listening success. To illustrate, one standard deviation change in modularity of the auditory-control network increased a listener’s chance of successful performance in a cued trial by about 3% on average, and it made her response about 40 ms faster. While the effect of modularity on behavior was found in states of both divided and selective attention, it was stronger for selective attention trials.

### Optimal brain network configuration for successful listening

The functional connectome was reconfigured towards more localized cortical processing during the listening task, as reflected by higher segregation of the whole-brain network. This functional specialization suggests that resolving the listening task mainly required attending to and processing the sentences as unconnected speech, arguably at the level of single words, which is often associated with cortical regions localized to auditory, cingulo-opercular, and inferior frontal regions under adverse listening situations (Obleser et al., 2007; DeWitt and Rauschecker, 2012; Vaden et al., 2013).

However, segregated topology is not neccessarily the optimal brain network configuration to resolve every difficult speech comprehension task. Instead, we argue that the most favourable configuration depends on the nature and instruction of the task, in particular on the level of linguistic and semantic information that need to be integrated to resolve the task (cf.Peelle, 2012; Lerner et al., 2011; de Heer et al., 2017). Indeed, previous studies in the contexts of working memory and complex reasoning have found a shift of the functional connectome towards higher network integration (Cohen and D’Esposito, 2016; Hearne et al., 2017). Along these lines of research, our findings emphasize the importance of the degree (rather than the direction) of modular reconfiguration of brain networks in predicting individuals’ behavior.

### Functional segregation within the auditory-control network is critical for cued speech comprehension

In adaptation to the listening task, a segregated module emerged along bilateral STS and STG with a posterior extension (Figure 4A/C, cortical maps; Figure 5C, dark blue module). This reconfiguration is in line with the proposal of a ventral stream in the functional neuroanatomy of speech processing (also referred to as the auditory ‘what’ stream) which has been suggested to be involved in sound-to-meaning mapping during speech recognition tasks (Alain et al., 2001; Ahveninen et al., 2006; Hickok and Poeppel, 2007; Saur et al., 2008; Rauschecker and Scott, 2009). In addition, the functional convergence of the auditory nodes with the posterior ventral attention nodes parallels the proposal that the posterior portion of the primary auditory cortex is involved in spatial separation of speech and sounds (Bushara et al., 1999; Zatorre et al., 2002; Arnott et al., 2004; Hill and Miller, 2010; Puschmann et al., 2017).

Our connectomic approach thus extends the existing evidence by demonstrating an increased functional cross-talk between neural systems involved in speech processing and sound localization during a cued speech comprehension task. Notably, the functional significance of this task-driven network reconfiguration was corroborated by its impact on listening success, in particular in states of selective attention. During selective attention trials, we assume an active attentional filter mechanism is being invoked to "tune out" the distracting sentence presented to the opposite ear, and to selectively amplify the sentence presented to the cued ear. Thus, the stronger brain-behavior correlation under selective attention trials suggests that this flexible filter is implemented by the formation of a more segregated and specialized cortical module composed of auditory and ventral attention nodes.

Another key observation here were auditory regions in anterior and superior temporal cortices as well as anterior insula and cingulate cortex merging with the somatomotor module during the listening task (Figure 4C, light blue). In parallel, the frontal ventral attention nodes in the vicinity of the IFG remained within the default-mode module (Figure 4C, dark blue). Within the auditory-control network, these reconfigurations appeared as the formation of highly segregated modules (Figure 5C, modules except the dark blue one). Importantly, the degree of the modular reconfiguration of cingulo-opercular and ventral attention nodes was predictive of listening success only when their network interactions were modelled in combination with the auditory nodes (i.e., the auditory-control network), and not with somatomotor or default-mode nodes.

Our findings concur conceptually with the earlier work by Vaden et al. (2013) who showed that, during a word recognition experiment, regions within the cingulo-opercular network displayed elevated activation in response to speech in noise, and regions along the ventral attention network showed this pattern during transitions between trials. Moreover, the authors found that cingulo-opercular regions exhibited higher functional connectivity relative to silence periods. Our results refine their findings and provide a clear distinction between cingulo-opercular and ventral attention nodes in their communication with auditory nodes during cued speech processing (Figure 5B). Our results support the hypothesis that, during a listening challenge, cingulo-opercular regions form an autonomous "core" control system that monitors effortful listening performance (Dosenbach et al., 2006; Vaden et al., 2015; Sadaghiani and D’Esposito, 2015), while ventral attention nodes implement attentional filtering of the relevant speech (Eckert et al., 2009).

Our connectomic approach can be distinguished from previous activation studies in two aspects. Firstly, we view the neural systems involved in challenging speech comprehension as a large-scale dynamic network whose configuration changes from "idling" or resting-state baseline to a high-duty, listening-task state. This allowed us to precisely delineate the cortical network embodiment of behavioral adaptation to a listening challenge.

Secondly, our graph-theoretical module identification unveiled a more complete picture (i.e., nodes, connections, and functional boundaries) of cortical subsystems supporting listening success. This allowed us to identify and characterize an auditory-control network whose modular organization revealed how functional coordination within and between auditory, cingulo-opercular, and ventral attention regions controls the selection and processing of speech during a listening challenge.

### Is modularity the network principle of auditory spatial attention?

The main finding of the current study is that higher modularity of the auditory-control network relative to resting state predicts individual listening success, both in states of divided and selective attention. Among the three network metrics, only modularity linked the reconfiguration of brain networks with this behavioral utilization of a spatial cue. This suggests that auditory spatial attention in adverse listening situations is implemented across cortex by a fine-tuned communication within and between neural assemblies across auditory, ventral attention and cingulo-opercular regions. This is in line with recent studies showing that higher modularity of brain networks supports auditory perception or decision-making (Sadaghiani et al., 2015; Alavash et al., 2017).

Combined with the principle of hierarchy (Simon, 1962; Kwapień and Drożdż, 2012), modularity allows for a formation of complex architectures that facilitate functional specificity, robustness and behavioral adaptation (Bassett et al., 2010; Bassett and Gazzaniga, 2011). Our results suggest that individuals’ ability in modifying intrinsic brain network modules is crucial for their attention to speech, and ultimately successful adaptation to a listening challenge. Notably, our results help better understand individual differences in speech comprehension at the systems level, with implications for hearing assistive devices and rehabilitation strategies.

### Limitations

The logic of modular reconfiguration of an auditory-control brain network in individual listeners implies it is possible to get reliable individual-specific estimates of network modularity. Recently, Gordon et al. (2017) demonstrated that reliability of functional connectivity differs greatly across individuals, measures of interest, and anatomical locations. Important to the present study, they showed that network modularity achieved an average difference of <3% from the split-data sample with only 10 min of data, while other network measures required longer scan duration. It is possible that the relatively short duration of the resting state scan in the present study may be hiding some significant individual variability. Future studies will be important to recover such important variation by dense sampling of individual brains (Poldrack, 2017; Satterthwaite et al., 2018).

Knowing that fMRI provides an indirect measure of neural activations, it remains unknown in how far our findings generalize to networks constructed based on brain neurophysiological responses recorded during resting state and listening (Keitel and Gross, 2016). In this respect, an important question is how network coupling between neural oscillations relates to listening behavior (Peelle and Davis, 2012; Giraud and Poeppel, 2012). In addition, although the sentences used in our linguistic Posner task were carefully designed to experimentally control individuals’ use of spatial and semantic cues during listening, their trial-by-trial presentation makes the listening task less naturalistic. Future studies will be important to investigate the brain network correlates of listening success during attention to continuous speech. Lastly, in the present study the brain networks were constructed based on correlations between regional time series which disregards directionality in the functional influences that regions may have on one another. Therefore, investigating causal relationships across cortical regions within the auditory-control network would help better understand the (bottom-up versus top-down) functional cross-talks between auditory, ventral attention and cingulo-opercular regions during challenging speech comprehension.

### Conclusion

In sum, our findings suggest that behavioral adaptation of a listener hinges on task-specific reconfiguration of the functional connectome at the level of network modules. We conclude that the communication dynamics within a fronto-temporal auditory-control network are closely linked with the successful deployment of auditory spatial attention in challenging listening situations.

## Materials and Methods

### Participants

Seventy-one healthy adult participants were invited from a larger cohort in which we investigate the neurocognitive mechanisms underlying adaptive listening in middle-aged and older adults (a large-scale study entitled "The listening challenge: How ageing brain adapt (AUDADAPT)"; https://cordis.europa.eu/project/rcn/197855_en.html).All participants were right-handed (assessed by a translated version of the Edinburgh Handedness Inventory (Oldfield, 1971) and had normal or corrected-to-normal vision. None of the participants had any history of neurological or psychiatric disorders. However, three participants had to be excluded following an incidental diagnosis made by an in-house radiologist based on the acquired structural MRI. In addition, fifteen participants were excluded because of incomplete measurements due to technical issues, claustrophobia, or task performance below chance level. Four additional participants completed the experiment but were removed from data analysis due to excessive head motion inside the scanner (i.e., scan-to-scan movement> 1.5 mm of translation or 1.5 degrees of rotation; root mean square (RMS) of frame-wise displacement (FD) > 0.5).

Accordingly, 49 participants were included in the main analysis (mean age 45.6 years, age range 19-69 years; 37 females). All procedures were in accordance with the Declaration of Helsinki and approved by the local ethics committee of the University of Lübeck. All participants gave written informed consent, and were financially compensated (8€ per hour).

### Procedure

Each imaging session consisted of seven fMRI runs (Figure 2A): resting state (∼8 min) followed by six runs in which participants performed a challenging speech comprehension task (∼9 min each; see Supplemental Information for detailed description of the task). Before the fMRI measurement, participants completed a short practice session of the listening task outside the scanner room to make sure they understood the task. Next, participants were placed in the scanner and went through a short sound volume adjustment to ensure balanced hearing between left and right ears at a comfortable level.

During the resting state scan participants were instructed to lie still inside the scanner, keep their eyes open and viewed a fixation cross displayed at the center of the screen, and to let their mind wander. A structural scan (∼5 min) was acquired at the end of the imaging session.

On a separate day before the imaging session, the older participants (>35 yrs) underwent a general screening procedure, detailed audiometry, and a battery of cognitive tests and personality profiling (see Tune et al. (2018) for details). Only participants with normal hearing or age-adequate mild-to-moderate hearing loss were invited for the imaging session.

### MRI data acquisition and preprocessing

Functional MRI data were collected by means of a Siemens MAGNETOM Skyra 3T scanner using a 64- channel head/neck coil and an echo-planar image (EPI) sequence [repetition time (TR) = 1000 ms; echo time (TE) = 25 ms; flip angle (FA) = 60°; acquisition matrix= 64 × 64; field of view (FOV) = 192 mm × 192 mm; voxel size= 3 × 3 × 3 mm; slice spacing= 3 mm]. Each image volume had 36 oblique axial slices parallel to the anterior commissure-posterior commissure (AC-PC) line, and was acquired with an acceleration factor of 2. Structural images were collected using a magnetization prepared rapid gradient echo (MP-RAGE) sequence [TR= 1900 ms; TE= 2.44 ms; FA= 9°; 1-mm isotropic voxel; 192 sagittal slices]. Although the acoustic noise associated with continuous imaging makes an auditory task more difficult, we used the same sequences for resting state and task with a comparable background scanner noise, which would cancel out in the task versus rest contrast.

#### Preprocessing

During the resting state 480 functional volumes were acquired, and during each run of the listening task 540 volume. To allow signal equilibration the first ten volumes of resting state and each run of the task were removed, and the remaining volumes were used for the subsequent analyses. Preprocessing steps were undertaken in SPM12. First, the functional images were spatially realigned to correct for head motion using a least square approach and the six rigid body affine transformations. Then, the functional volumes were corrected for slice timing to adjust for differences in image acquisition times between slices. Subsequently, functional images were coregistered to each individual’s structural image. Next, the normalization parameters from the subject’s native space to MNI space were obtained via the unified segmentation of their Tl image (Ashburner and Friston, 2005). The normalization parameters (warps) were then applied to each of the EPI volumes (Ashburner and Friston, 2005). The resulting functional volumes and the structural volume were then spatially normalized to the SPM single subject Tl volume in MNI space. No spatial smoothing was applied on the volumes to avoid potential artificial distance-dependent correlations between voxels’ BOLD signals (Fornito et al., 2013; Stanley et al., 2013).

#### Motion artifact removal

In recent years, the potentially confounding impact of head motion or non-neural physiological trends on the temporal correlations between BOLD signals has been raised by multiple studies (see Murphy et al., 2013; Power et al, 2015 for reviews). Knowing that older participants often make larger within-scan movements than younger participants (Van Dijk et al., 2012; Geerligs et al., 2015) the issue of motion is particularly important in the current study where the age of the participants spans across a wide range. Thus, we accounted for motion artifacts in three stages of our analysis (Figure 1B). First, we used the denoising method proposed by (Patel et al., 2014) that decomposes brain signals in wavelet space, identifies and removes trains of spikes that likely reflect motion artifacts, and reconstitutes denoised signals from the remaining wavelets. Second, we implemented a nuisance regression to suppress non-neural and motion artifacts as explained in the next section. Finally, we used RMS of frame-wise displacement as a regressor of no-interest in the mixed-effects analyses (see *Statistical analysis).*

#### Cortical parcellation

Cortical nodes were defined using a previously established functional parcellation (Gordon et al., 2016) which has shown relatively higher accuracy as compared to other parcellations (Arslan et al., 2017). This parcellation is based on surface-based resting state functional connectivity boundary maps and encompasses 333 cortical areas. We used this template to estimate mean BOLD signals across voxels within each cortical region per participant (Figure 1B).

#### Nuisance regression

To minimize the effects of spurious temporal correlations induced by physiological and movement artifacts, a general linear model (GLM) was constructed to regress out white matter and cerebrospinal fluid (CSF) mean time series, together with the six rigid-body movement parameters and their first-order temporal derivatives (Hallquist et al., 2013). Subsequently, the residual time series obtained from this procedure were concatenated across the six runs of the listening task for each participant. The cortical time series of the resting state and listening task were further used for functional connectivity and network analysis (Figure 1B). Note that global signal was not regressed out from the regional time series, as it is still an open question in the field what global signal regression (GSR) in fact regresses out. On the one hand, it has been shown that large amount of the global signal variance can be attributed to respiration, head motion, and scanner hardware-related artifacts, and that global signal has mainly a sensorimotor disruption map (Power et al., 2017). On the other hand, the global signal has also been shown to be tightly coupled with underlying neural signals (Schölvinck et al., 2010). Further, doing GSR shifts the distribution of functional connectivity values from being predominantly positive to being centered around zero, with the meaning of the negative correlations being unclear (Murphy and Fox, 2016). In the present study, wavelet despiking of motion artefacts allowed us to accommodate the spatial and temporal heterogeneity of motion artifacts. Importantly, our main results point to the reconfiguration of a fronto-temporal auditory-control network, which does not overlap with the mainly sensorimotor disruption map of the global signal.

### Functional connectivity

First, mean residual time series were band-pass filtered by means of maximum overlap discrete wavelet transform (Daubechies wavelet of length 8), and the results of the filtering within the range of 0.06-0.12 Hz (wavelet scale 3) were used for further analyses. It has been previously documented that the behavioral correlates of the functional connectome are best observed by analyzing low-frequency large-scale brain networks (Salvador et al., 2005; Achard et al., 2006; Achard et al., 2008; Giessing et al., 2013; Alavash et al., 2015b). The use of wavelet scale 3 was motivated by previous work showing that behavior during cognitive tasks predominately correlated with changes in functional connectivity in the same frequency range (Bassett et al., 2010; Alavash et al., 2015a; Alavash et al., 2016; Alavash et al., 2018). To obtain a measure of association between each pair of cortical regions, Fisher’s Z-transformed Pearson correlations between wavelet coefficients were computed, which resulted in one 333 × 333 correlation matrix per participant for each resting state and listening task (Figure 1B).

### Connectomics

Brain graphs were constructed from the functional connectivity matrices by including the top 10% of the connections in the graph according to the rank of their correlation strengths (Ginestet et al., 2011; Fornito et al., 2013). This resulted in sparse binary undirected brain graphs at a fixed network density of 10%, and assured that the brain graphs were matched in terms of density across participants, resting state and listening task (van Wijk et al., 2010; van den Heuvel et al., 2017). To investigate whether choosing a different (range of) graph density threshold(s) would affect the results (Garrison et al., 2015), we examined the effect of graph thresholding using the cost-integration approach (Ginestet et al., 2014; Figure S1).

Mean functional connectivity was calculated as the average of the upper-diagonal elements of the sparse connectivity matrix for each participant. In addition, three key topological network metrics were estimated: mean local efficiency, network modularity, and global network efficiency. These graph-theoretical metrics were used to quantify the configuration of large-scale brain networks on the local, intermediate, and global scales of topology, respectively (Figure 1A; Betzel and Bassett, 2016). For each topological metric, we computed a whole-brain metric which collapses that network property into one single metric (Brain Connectivity Toolbox; Rubinov and Sporns, 2010). The mathematical formalization of these measures are provided in the Supplementary Information. Mean within- (between-) module connectivity was estimated as the sum of connection weights falling within (between) modules, normalized by maximum number of possible within- (between) connections, respectively (Garcia et al., 2018).

### Statistical analysis

#### Behavioral data

Performance of the participants during the listening task was measured based on (a binary measure of) correct identification of the probed sentence-final word (accuracy) and the inverse of reaction time on correct trials (response speed). Trials on which no response was given (within the response time window) were considered as incorrect. The single-trial behavioral measures across all individual participants were treated as the dependent variable in the (generalized) linear mixed-effects analysis (see *Statistical analysis).*

#### Brain data

Statistical comparisons of network metrics between resting state and listening task were based on permutation tests for paired samples (randomly permuting the rest and task labels 10,000 times). We used Cohen’s d for paired samples as the corresponding effect size (Hentschke and Stuttgen, 2011).

#### Generalized linear mixed-effects models

Brain-behavior relationship was investigated within a linear mixed-effects analysis framework To this end, either of the single-trial behavioral measures (accuracy or response speed) across all participants were treated as the dependent variable. The main predictors in the model were the spatial and semantic cues, each at two levels (divided vs. selective and general vs. specific, respectively). Since in the current study the main question is whether reconfiguration of resting state brain networks in adaptation to the listening challenge relates to listening success, we used the difference between a given network parameter across resting state and the listening task (i.e., task minus rest) as the third main predictor in the model. The linear mixed-effects analysis framework allowed us to control for other variables which entered as regressors of no-interest in the model. These included age, head motion (i.e., difference in RMS of frame-wise displacement between resting state and task) and side probed (left or right). Mixed-effects analyses were performed in R (R Core Team, 2017) using the packages lme4 (Bates et al., 2015) and sjPlot (Lüdecke, 2018).

#### Model selection

To estimate the best fit of the linear mixed-effects models, we followed an iterative model fitting procedure, which started with an intercept-only null model (Tune et al., 2018). Fixed effects terms were added in a stepwise procedure, and the change of model fit (performed using maximum-likelihood estimation) was assessed using likelihood ratio tests. Deviation coding was used for categorical predictors. All continuous variables were Z-scored. Since accuracy is a binary variable, we used generalized linear mixed-effects model with the logit as link function. In the case of response speed, the distribution of the data did not significantly deviate from normal distribution (Shapiro-Wilk test p-value = 0.7). Thus, in this case we used linear mixed-effects models with the underlying distribution set to Gaussian. For both models predicting accuracy or response speed we report P-values for individual model terms that were derived using the Satterthwaite approximation for degrees of freedom (Luke, 2017). As a measure of effects size, for the model predicting accuracy we report odds ratios (OR) and for response speed we report the regression coefficient (β).

### Data visualization

Brain surfaces were visualized using the Connectome Workbench v.1.2.3. Connectograms were visualized using the Brain Data Viewer application. Network flow diagrams were visualized in MapEquation application.

### Data availability

The complete dataset associated with this work including raw data and connectivity maps as well as code of network and statistical analyses is publicly available under https://osf.io/28r57/.

## Acknowledgement

Research was supported by the European Research Council (ERC Consolidator grant AUDADAPT, no. 646696) to JO. Anne Herrmann, Susanne Schellbach, Franziska Scharata, and Leonhard Waschke helped acquire, manage, and archive the data.

## References

Achard S, Bassett DS, Meyer-Lindenberg A, Bullmore E (2008) Fractal connectivity of long-memory networks. Phys Rev E 77.

Achard S, Salvador R, Whitcher B, Suckling J, Bullmore E (2006) A resilient, low-frequency, small-world human brain functional network with highly connected association cortical hubs. The Journal of neuroscience: the official journal of the Society for Neuroscience 26:63–72.

Adank P (2012) The neural bases of difficult speech comprehension and speech production: Two Activation Likelihood Estimation (ALE) meta-analyses. Brain Lang 122:42–54.

Ahveninen J, Jaaskelainen IP, Raij T, Bonmassar G, Devore S, Hamalainen M, Levanen S, Lin FH, Sams M, Shinn-Cunningham BG, Witzel T, Belliveau JW (2006) Task-modulated "what" and "where" pathways in human auditory cortex. Proc Natl Acad Sci USA 103:14608–14613.

Alain C, Arnott SR, Hevenor S, Graham S, Grady CL (2001) "What" and "where" in the human auditory system. Proc Natl Acad Sci USA 98:12301–12306.

Alavash M, Thiel CM, Giessing C (2016) Dynamic coupling of complex brain networks and dual-task behavior. Neurolmage 129:233–246.

Alavash M, Hilgetag CC, Thiel CM, Giessing C (2015a) Persistency and flexibility of complex brain networks underlie dual-task interference. Hum Brain Mapp 36:3542–3562.

Alavash M, Doebler P, Holling H, Thiel CM, Giessing C (2015b) Is functional integration of resting state brain networks an unspecific biomarker for working memory performance? Neurolmage 108:182–193.

Alavash M, Daube C, Wöstmann M, Brandmeyer A, Obleser J (2017) Large–scale network dynamics of beta–band oscillations underlie auditory perceptual decision–making. Network Neuroscience 1:166–191.

Alavash M, Lim SJ, Thiel C, Sehm B, Deserno L, Obleser J (2018) Dopaminergic modulation of hemodynamic signal variability and the functional connectome during cognitive performance. Neurolmage 172:341–356.

Arnott SR, Binns MA, Grady CL, Alain C (2004) Assessing the auditory dual-pathway model in humans. Neurolmage 22:401–408.

Arslan S, Ktena SI, Makropoulos A, Robinson EC, Rueckert D, Parisot S (2017) Human brain mapping: A systematic comparison of parcellation methods for the human cerebral cortex. Neurolmage 170:5–30.

Ashburner J, Friston KJ (2005) Unified segmentation. Neurolmage 26:839–851.

Bassett DS, Gazzaniga MS (2011) Understanding complexity in the human brain. Trends Cogn Sci 15:200–209.

Bassett DS, Wymbs NF, Porter MA, Mucha PJ, Carlson JM, Grafton ST (2010) Dynamic reconfiguration of human brain networks during learning. Proc Natl Acad Sci USA 108:7641–7646.

Bates D, Mächler M, Bolker B, Walker S (2015) Fitting linear mixed-effects models using lme4. Journal of Statistical Software 67.

Betzel RF, Bassett DS (2016) Multi-scale brain networks. Neurolmage 160:73–83.

Blondel S, Guillaume J, Lambiotte R, Lefebvre E (2008) Fast unfolding of communities in large networks. J Stat Mech P10008:6.

Bolt T, Nomi JS, Rubinov M, Uddin LQ (2017) Correspondence between evoked and intrinsic functional brain network configurations. Hum Brain Mapp.

Bullmore ET, Bassett DS (2011) Brain graphs: graphical models of the human brain connectome. Annu RevClin Psychol 7:113–140.

Bullmore E, Sporns O (2009) Complex brain networks: graph theoretical analysis of structural and functional systems. Nature Rev Neurosci 10:186–198.

Bushara K, Weeks R, Ishii K, Catalan M, Tian B, Rauschecker J, Hallett M (1999) Modality-specific frontal and parietal areas for auditory and visual spatial localization in humans. Nature Neurosci 2:759–766.

Corbetta M, Patel G, Shulman GL (2008) The reorienting system of the human brain: from environment to theory of mind. Neuron 58:306–324.

Chai LR, Mattar MG, Blank IA, Fedorenko E, Bassett DS (2016) Functional network dynamics of the language system. Cereb Cortex 26:4148–4159.

Cohen JR, D’Esposito M (2016) The Segregation and Integration of Distinct Brain Networks and Their Relationship to Cognition. J Neurosci 36:12083–12094.

Cole MW, Bassett DS, Power JD, Braver TS, Petersen SE (2014) Intrinsic and task-evoked network architectures of the human brain. Neuron 83:238–251.

Colflesh G, Conway A (2007) Individual differences in working memory capacity and divided attention in dichotic listening. Psychon Bull Rev 14:699–703.

de Heer WA, Huth AG, Griffiths TL, Gallant JL, Theunissen FE (2017) The hierarchical cortical organization of human speech processing. J Neurosci 37:6539–6557.

Dai L, Best V, Shinn-Cunningham BG (2018) Sensorineural hearing loss degrades behavioral and physiological measures of human spatial selective auditory attention. Proc Natl Acad Sci USA 115:E3286–E3295.

DeWitt I, Rauschecker JP (2012) Phoneme and word recognition in the auditory ventral stream. Proc Natl Acad Sci USA 109:E505–514.

Dosenbach NU, Fair DA, Miezin FM, Cohen AL, Wenger KK, Dosenbach RA, Fox MD, Snyder AZ, Vincent JL, Raichle ME, Schlaggar BL, Petersen SE (2007) Distinct brain networks for adaptive and stable task control in humans. Proc Natl Acad Sci USA 104:11073–11078.

Dosenbach NU, Visscher KM, Palmer ED, Miezin FM, Wenger KK, Kang HC, Burgund ED, Grimes AL, Schlaggar BL, Petersen SE (2006) A core system for the implementation of task sets. Neuron 50:799–812.

Eckert MA, Menon V, Walczak A, Ahlstrom J, Denslow S, Horwitz A, Dubno JR (2009) At the heart of the ventral attention system: the right anterior insula. Hum Brain Mapp 30:2530–2541.

Erb J, Henry MJ, Eisner F, Obleser J (2013) The brain dynamics of rapid perceptual adaptation to adverse listening conditions. J Neurosci 33:10688–10697.

Fedorenko E, Thompson-Schill S (2014) Reworking the language network. Trends Cogn Sci 18:120–126.

Finn ES, Shen X, Scheinost D, Rosenberg MD, Huang J, Chun MM, Papademetris X, Constable RT (2015) Functional connectome fingerprinting: identifying individuals using patterns of brain connectivity. Nature Neurosci 18:1664–1671.

Fornito A, Zalesky A, Breakspear M (2013) Graph analysis of the human connectome: promise, progress, and pitfalls. Neurolmage 80:426–444.

Fuertinger S, Horwitz B, Simonyan K (2015) The Functional Connectome of Speech Control. PLoS Biol 13:e1002209.

Garcia JO, Ashourvan A, Muldoon SF, Vettel JM, Bassett DS (2018) Applications of community detection techniques to brain graphs: Algorithmic considerations and implications for neural function. Proceedings of the IEEE:1–22.

Garrison KA, Scheinost D, Finn ES, Shen X, Constable RT (2015) The (in)stability of functional brain network measures across thresholds. Neurolmage 118:651–661.

Geerligs L, Rubinov M, Cam C, Henson RN (2015) State and Trait Components of Functional Connectivity: Individual Differences Vary with Mental State. J Neurosci 35:13949–13961.

Giessing C, Thiel CM, Alexander-Bloch AF, Patel AX, Bullmore ET (2013) Human brain functional network changes associated with enhanced and impaired attentional task performance. J Neurosci 33:5903–5914.

Ginestet CE, Nichols TE, Bullmore ET, Simmons A (2011) Brain network analysis: separating cost from topology using cost-integration. PloS one 6:e21570.

Giraud AL, Poeppel D (2012) Cortical oscillations and speech processing: emerging computational principles and operations. Nature Neurosci 15:511–517.

Gordon EM, Laumann TO, Gilmore AW, Newbold DJ, Greene DJ, Berg JJ, Ortega M, Hoyt-Drazen C, Gratton C, Sun H, Hampton JM, Coalson RS, Nguyen AL, McDermott KB, Shimony JS, Snyder AZ, Schlaggar BL, Petersen SE, Nelson SM, Dosenbach NUF (2017) Precision functional mapping of individual human brains. Neuron 95:791–807 e797.

Gordon EM, Laumann TO, Adeyemo B, Huckins JF, Kelley WM, Petersen SE (2016) Generation and evaluation of a cortical area parcellation from resting-state correlations. Cereb Cortex 26:288–303.

Gordon EM, Laumann TO, Adeyemo B, Petersen SE (2017a) Individual Variability of the System-Level Organization of the Human Brain. Cereb Cortex 27:386–399.

Gordon EM, Laumann TO, Adeyemo B, Gilmore AW, Nelson SM, Dosenbach NUF, Petersen SE (2017b) Individual-specific features of brain systems identified with resting state functional correlations. Neurolmage 146:918–939.

Gratton C, Laumann TO, Gordon EM, Adeyemo B, Petersen SE (2016) Evidence for Two Independent Factors that Modify Brain Networks to Meet Task Goals. Cell Rep 17:1276–1288.

Gratton C, Laumann TO, Nielsen AN, Greene DJ, Gordon EM, Gilmore AW, Nelson SM, Coalson RS, Snyder AZ, Schlaggar BL, Dosenbach NUF, Petersen SE (2018) Functional Brain Networks Are Dominated by Stable Group and Individual Factors, Not Cognitive or Daily Variation. Neuron 98:439–452 e435.

Hagoort P (2014) Nodes and networks in the neural architecture for language: Broca’s region and beyond. Curr Opin Neurobiol 28:136–141.

Hallquist MN, Hwang K, Luna B (2013) The nuisance of nuisance regression: spectral misspecification in a common approach to resting-state fMRI preprocessing reintroduces noise and obscures functional connectivity. Neurolmage 82:208–225.

Hearne LJ, Cocchi L, Zalesky A, Mattingley JB (2017) Reconfiguration of brain network architectures between resting state and complexity-dependent cognitive reasoning. J Neurosci.

Hentschke H, Stuttgen MC (2011) Computation of measures of effect size for neuroscience data sets. Eur J Neurosci 34:1887–1894.

Hickok G, Poeppel D (2007) The cortical organization of speech processing. Nature Rev Neurosci 8 393–402.

Hill KT, Miller LM (2010) Auditory attentional control and selection during cocktail party listening. Cereb Cortex 20:583–590.

Honey CJ, Kotter R, Breakspear M, Sporns O (2007) Network structure of cerebral cortex shapes functional connectivity on multiple time scales. Proc Natl Acad Sci USA 104:10240–10245.

Hugdahl, K. (2003). Dichotic Listening in the study of auditory laterality. In K. Hugdahl & R. J. Davidson (Eds.), The asymmetrical brain (pp. 441–475). Cambridge, MA, US: MIT Press.

Keitel A, Gross J (2016) Individual human brain areas can be identified from their characteristic spectral activation fingerprints. PLOS Biology 14:e1002498.

Keitel A, Ince RAA, Gross J, Kayser C (2017) Auditory cortical delta-entrainment interacts with oscillatory power in multiple fronto-parietal networks. Neurolmage 147:32–42.

Kidd GR, Watson CS, Gygi B (2007) Individual differences in auditory abilities. J Acoust Soc Am 122:418–435.

Kwapień J, Drożdż S (2012) Physical approach to complex systems. Physics Reports 515:115–226.

Laumann TO, Snyder AZ, Mitra A, Gordon EM, Gratton C, Adeyemo B, Gilmore AW, Nelson SM, Berg JJ, Greene DJ, McCarthy JE, Tagliazucchi E, Laufs H, Schlaggar BL, Dosenbach NUF, Petersen SE (2017) On the Stability of BOLD fMRI Correlations. Cereb Cortex 27:4719–4732.

Lerner Y, Honey CJ, Silbert LJ, Hasson U (2011) Topographic mapping of a hierarchy of temporal receptive windows using a narrated story. J Neurosci 31:2906–2915.

Lüdecke D (2018) sjPlot: Data visualization for statistics in social science. R package version 2.4.1, https://CRAN.R-project.org/package=sjPlot.

Mueller S, Wang D, Fox MD, Yeo BT, Sepulcre J, Sabuncu MR, Shafee R, Lu J, Liu H (2013) Individual variability in functional connectivity architecture of the human brain. Neuron 77:586–595.

Muller AM, Meyer M (2014) Language in the brain at rest: new insights from resting state data and graph theoretical analysis. Front Hum Neurosci 8:228.

Murphy K, Fox MD (2016) Towards a consensus regarding global signal regression for resting state functional connectivity MRI. Neurolmage.

Murphy K, Birn RM, Bandettini PA (2013) Resting-state fMRI confounds and cleanup. Neurolmage 80:349–359.

Obleser J, Kotz SA (2010) Expectancy constraints in degraded speech modulate the language comprehension network. Cereb Cortex 20:633–640.

Obleser J, Wise RJ, Dresner MA, Scott SK (2007) Functional integration across brain regions improves speech perception under adverse listening conditions. J Neurosci 27:2283–2289.

Oldfield, R.C. (1971) The assessment and analysis of handedness: the Edinburgh inventory. Neuropsychologia, 9, 97–113.

Passow S, Muller M, Westerhausen R, Hugdahl K, Wartenburger I, Heekeren HR, Lindenberger U, Li SC (2013) Development of attentional control of verbal auditory perception from middle to late childhood: comparisons to healthy aging. Dev Psychol 49:1982–1993.

Patel AX, Kundu P, Rubinov M, Jones PS, Vertes PE, Ersche KD, Suckling J, Bullmore ET (2014) A wavelet method for modeling and despiking motion artifacts from resting-state fMRI time series. Neurolmage 95:287–304.

Peelle JE (2017) Listening Effort: How the Cognitive Consequences of Acoustic Challenge Are Reflected in Brain and Behavior. Ear Hear.

Peelle JE (2012) The hemispheric lateralization of speech processing depends on what "speech" is: a hierarchical perspective. Front Hum Neurosci 6:309.

Peelle JE, Davis MH (2012) Neural Oscillations Carry Speech Rhythm through to Comprehension. Front Psychol 3:320.

Power JD, Cohen AL, Nelson SM, Wig GS, Barnes KA, Church JA, Vogel AC, Laumann TO, Miezin FM, Schlaggar BL, Petersen SE (2011) Functional network organization of the human brain. Neuron 72:665–678.

Power JD, Plitt M, Laumann TO, Martin A (2017) Sources and implications of whole-brain fMRI signals in humans. Neurolmage 146:609–625.

Power JD, Schlaggar BL, Petersen SE (2015) Recent progress and outstanding issues in motion correction in resting state fMRI. Neurolmage 105C:536–551.

Poldrack RA (2017) Precision neuroscience: Dense sampling of individual brains. Neuron 95:727–729.

Puschmann S, Steinkamp S, Gillich I,Mirkovic B, Debener S, Thiel CM (2017) The right temporoparietal junction supports speech tracking during selective listening: Evidence from concurrent EEG-fMRI. J Neurosci 37:11505–11516.

Price C, Thierry G, Griffiths T (2005) Speech-specific auditory processing: where is it? Trends in cognitive sciences 9:271–276.

Rauschecker JP, Scott SK (2009) Maps and streams in the auditory cortex: nonhuman primates illuminate human speech processing. Nature Neurosci 12:718–724.

Raichle ME, Macleod AM, Snyder AZ, Powers WG, Gusnard DA, Shulman GL (2000) A default mode of brain function. Proc Natl Acad Sci USA 98:676–682.

Rubinov M, Sporns O (2010) Complex network measures of brain connectivity: uses and interpretations. Neurolmage 52:1059–1069.

Sadaghiani S, D’Esposito M (2015) Functional Characterization of the Cingulo-Opercular Network in the Maintenance of Tonic Alertness. Cereb Cortex 25:2763–2773.

Sadaghiani S, Poline JB, Kleinschmidt A, D’Esposito M (2015) Ongoing dynamics in large-scale functional connectivity predict perception. Proc Natl Acad Sci USA 112:8463–8468.

Salvador R, Suckling J, Coleman MR, Pickard JD, Menon D, Bullmore E (2005) Neurophysiological architecture of functional magnetic resonance images of human brain. Cereb Cortex 15:1332–1342.

Saur D, Kreher BW, Schnell S, Kummerer D, Kellmeyer P, Vry MS, Umarova R, Musso M, Glauche V, Abel S, Huber W, Rijntjes M, Hennig J, Weiller C (2008) Ventral and dorsal pathways for language. Proc Natl Acad Sci USA 105:18035–18040.

Schölvinck M, Maier A, Ye FQ, Duyn JH, Leopold DA (2010) Neural basis of global resting-state fMRI activity. PNAS 107(22): 10238–43.

Schultz DH, Cole DM (2016) Higher Intelligence Is Associated with Less Task-Related Brain Network Reconfiguration. J Neurosci 36:38551–38561.

Shen K, Hutchison RM, Bezgin G, Everling S, McIntosh AR (2015) Network structure shapes spontaneous functional connectivity dynamics. The Journal of neuroscience : the official journal of the Society for Neuroscience 35:5579–5588.

Simon H (1962) The architecture of complexity. Proc Am Phil Soc 106.

Simony E, Honey CJ, Chen J, Lositsky O, Yeshurun Y, Wiesel A, Hasson U (2016) Dynamic reconfiguration of the defauIt mode network during narrative comprehension. Nat Commun 7:12141.

Spadone S, Della Penna S, C S, Betti V, Tosoni A, Perrucci M, Romani G, Corbetta M (2015) Dynamic reorganization of human resting-state networks during visuospatial attention. Proc Natl Acad Sci USA 112:8112–8117.

Sporns O (2013) Network attributes for segregation and integration in the human brain. Cur Opin Neurobiol 23:162–171.

Stanley ML, Moussa MN, Paolini BM, Lyday RG, Burdette JH, Laurienti PJ (2013) Defining nodes in complex brain networks. Front Comput Neurosci 7:169.

Satterthwaite TD, Xia CH, Bassett DS (2018) Personalized neuroscience: Common and individual-specific features in functional brain networks. Neuron 98:243–245.

Tavor I, Parker Jones O, Mars RB, Smith SM, Behrens TE, Jbabdi S (2016) Task-free MRI predicts individual differences in brain activity during task performance. Science 352:216–220.

Tomasi D, Volkow ND (2012) Resting functional connectivity of language networks: characterization and reproducibility. Mol Psychiatry 17:841–854.

Tune S, Wostmann M, Obleser J (2018) Probing the limits of alpha power lateralisation as a neural marker of selective attention in middle-aged and older listeners. Eur J Neurosci.

Vaden KI, Jr., Kuchinsky SE, Cute SL, Ahlstrom JB, Dubno JR, Eckert MA (2013) The cingulo-opercular network provides word-recognition benefit. J Neurosci 33:18979–18986.

Vaden KI, Jr., Kuchinsky SE, Ahlstrom JB, Dubno JR, Eckert MA (2015) Cortical activity predicts which older adults recognize speech in noise and when. J Neurosci 35:3929–3937.

van den Heuvel M, de Lange S, Zalesky A, Seguin C, Yeo T, Schmidt R (2017) Proportional thresholding in resting-state fMRI functional connectivity networks and consequences for patient-control connectome studies: Issues and recommendations. Neurolmage.

van Dijk KR, Sabuncu MR, Buckner RL (2012) The influence of head motion on intrinsic functional connectivity MRI. Neurolmage 59:431–438.

van Wijk BCM, Stam CJ, Daffertshofer A (2010) Comparing Brain Networks of Different Size and Connectivity Density Using Graph Theory. PloS one 5:e13701.

Wild CJ, Yusuf A, Wilson DE, Peelle JE, Davis MH, Johnsrude IS (2012) Effortful listening: the processing of degraded speech depends critically on attention. J Neurosci 32:14010–14021.

Wöstmann M, Herrmann B, Maess B, Obleser J (2016) Spatiotemporal dynamics of auditory attention synchronize with speech. Proc Natl Acad Sci USA 113:3873–3878.

Zatorre RJ, Bouffard M, Ahad P, Belin P (2002) Where is ‘where’ in the human auditory cortex? Nature Neurosci 5:905–909.

